# Balance of osmotic pressures determines the volume of the cell nucleus

**DOI:** 10.1101/2021.10.01.462771

**Authors:** Dan Deviri, Samuel A. Safran

**Affiliations:** Department of Chemical and Biological Physics, Weizmann Institute of Science, Rehovet 76100, Israel

## Abstract

The volume of the cell nucleus varies across cell-types and species, and is commonly thought to be determined by the size of the genome and degree of chromatin compaction. However, this notion has been challenged over the years by multiple experimental evidence. Here, we consider the physical condition of mechanical force balance as a determining condition of the nuclear volume and use quantitative, order-of-magnitude analysis to estimate the forces from different sources of nuclear and cellular pressure. Our estimates suggest that the dominant pressure within the nucleus and cytoplasm originates from the osmotic pressure of proteins and RNA molecules that are localized to the nucleus or cytoplasm by out-of-equilibrium, active nucleocytoplasmic transport rather than from chromatin or its associated ions. This motivates us to formulate a physical model for the ratio of the cell and nuclear volumes in which osmotic pressures of localized proteins determine the relative volumes. In accordance with unexplained observations that are century-old, our model predicts that the ratio of the cell and nuclear volumes is a constant, robust to a wide variety of biochemical and biophysical manipulations, and is changed only if gene expression or nucleocytoplasmic transport are modulated.

## 1 Introduction

The nucleus, which is the largest organelle in the cell, regulates the expression of genes [1] via transcription of DNA found in the chromatin (a complex of DNA polymer and histone proteins), protects genes from mechanical and biochemical damage [2, 3], and controls their environment [1]. Furthermore, nuclear size is related to chromatin organization [4], and changes of the nuclear size accompany cell differentiation, development, and disease [5]. However, despite the effect of nuclear volume changes on chromatin organization, the associated biophysical mechanisms that determine it are not well understood. Recently, Cantwell and Nurse [6] reviewed the biological factors that are implicated in nuclear size determination: cell size, cytoplasmic factors, the LINC complex, transcription and RNA processing, nucleocytoplasmic transport, and nuclear envelope (NE) expansion. The goal of the present work is to translate these biological factors into quantifiable physical quantities, in particular the force balance between the nucleus and cytoplasm that in mechanical (but not necessarily thermodynamic) equilibrium, determines nuclear volume. Our calculations show that the primary forces are due to the osmotic pressures of proteins and RNA molecules preferentially localized to the nucleus and cytoplasm and that this implies that under many conditions, the ratio of the nuclear and cell volumes is a constant – an observation that was first made over 100 years ago [7] and has been further quantified in more recent studies [6].

As noted in the biological review [6], several experiments suggest that the chromatin content of the nucleus does not determine its size, in both yeasts [8, 9], C. elegans [10], and vertebrates [11, 12]. This conclusion disagrees with the widely accepted, qualitative “*nucleoskeletal theory*” that hypothesizes that the degree of chromatin compaction by the cellular machinery is the determining factor of nuclear size. This implies that nuclear size should sensitively depend on chromatin content [13] which is assumed to completely fill the nuclear volume. This assumption is directly refuted by some imaging experiments that observed that the chromatin does not always fill the entire volume of the nucleus [2, 14, 15]. These experiments, together with others reviewed in [6], indicate that the popular “*nucleoskeletal theory*” does not adequately describe the biophysical mechanisms that underlie nuclear size determination.

In this paper, we present a comparison of the possible biophysical forces that may determine the volume of the nucleus. We relate the biological factors reviewed by Cantwell and Nurse, as well as the size of the genome, to six potentially significant physical forces that originate from different components of the nucleus and cytoplasm: (1) osmotic pressure of the chromatin polymers, (2) osmotic pressure of counterions that balance the chromatin charge, (3) elastic forces of the cytoskeleton, (4) osmotic pressures of solutes (proteins and RNA molecules) that are preferentially localized (due to active transport) to the cytoplasm or nucleoplasm, and (5) NE tension. Each of these forces is physically derived and quantified using published biological data. Our estimates suggest that in the common scenario where the NE is not stretched, the osmotic pressures of preferentially-localized, large molecule solutes (proteins, RNA) in the nucleus are the dominant forces whose balance with those in the cytoplasm determine the nuclear volume. Motivated by this result, we formulate a minimal biophysical model that relates both the volumes of the cell and the nucleus to mechanical equilibrium across the plasma membrane and NE. Our model predicts that for relaxed (i.e., not taut) NE, the volumes of the cell and the nucleus are proportional even when various biophysical factors (e.g. osmotic pressure, cellular adhesion, genome size) are varied. This result agrees with a biological observation that is more than a century old [6, 7], which underlies a popular, clinical screening techniques for cancer cells, for which the proportion between the cell and nuclear volumes deviates from specific values [16]. Besides suggesting a mechanistic explanation for the proportionality of cell and nuclear volumes, our physical theory highlights the potential importance of currently-underestimated effects of protein and RNA localization (via nucleocytoplasmic transport) in nuclear mechanics. Our results and predictions may impact several different fields, including medicine, cellular biology, and nuclear mechanobiology. We begin by reviewing the physical forces that influence the volume of the nucleus, and quantifying their effects using physical theory. The various estimates are compared and the conditions for which the cell and nuclear volumes are proportional are elucidated. Finally, we discuss the relation of the theory to various experiments.

## 2 Physical forces that may determine nuclear volume

There are two types of physical forces that deform bodies, in our case the cell nucleus: (i) Isotropic forces, known as pressures, which change the volume of the object but not its shape, and (ii) Directional, shear forces, which change the shape of the object but not its volume. Since we are interested in the volume of the nucleus and not in its detailed shape, we focus on the pressures; the effects of shear forces are discussed in the Discussion section. In the idealized case that there are no shear forces (e.g., due to stress fibers or contractility of neighboring cells [17]), and when the NE is homogeneous then the nucleus is spherical, which is indeed the case in many biological scenarios [18].

The nuclear volume is determined by the balance of inward pressures which compress the nucleus and usually originate from the cytoplasm, and outward pressures that expand the nucleus and usually originate from within the nucleoplasm (see Fig. 1). An imbalance of the inward and outward pressures changes the volume of the nucleus by facilitating flow of fluid either into or out of the nucleus through the nuclear pore complexes (NPCs) that are embedded in the NE. Mechanical stability dictates that the change in nuclear volume that follows the flow of solvent must decrease the imbalance between inward and outward pressures, eventually balancing the two. In steady state, the system is in mechanical equilibrium so that the inward pressures and outward pressures that originate from the cytoplasm and nucleoplasm are balanced by the inward pressure associated with the surface tension of the NE; this is formulated by Young-Laplace law (see Fig. 1 and SI of [19] for a derivation)

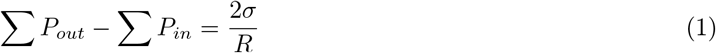

where Σ*P_out_* and Σ*P_in_* are respectively the sums of all the outward and inward pressures, *σ* is the NE tension, and *R* is the radius of the nucleus; in cases where the nucleus is not spherical, 2/*R* on the right hand side of Eq. 1 is replace by the sum of the local radii of curvatures, which may vary in space if the nuclear shape is not spherical. In the subsections below, we estimate the order of magnitude of each of these inward and outward pressures and relate them to biological structures and functions discussed in [6]. To demonstrate the generality of our estimates, we consider nuclei of two eukaryotic organisms from different biological kingdoms, humans (animalia) and the fission yeast S. Pombe (fungi); our estimates of the various forces in the two species are summarized in Table 1 below.

**Figure 1:**
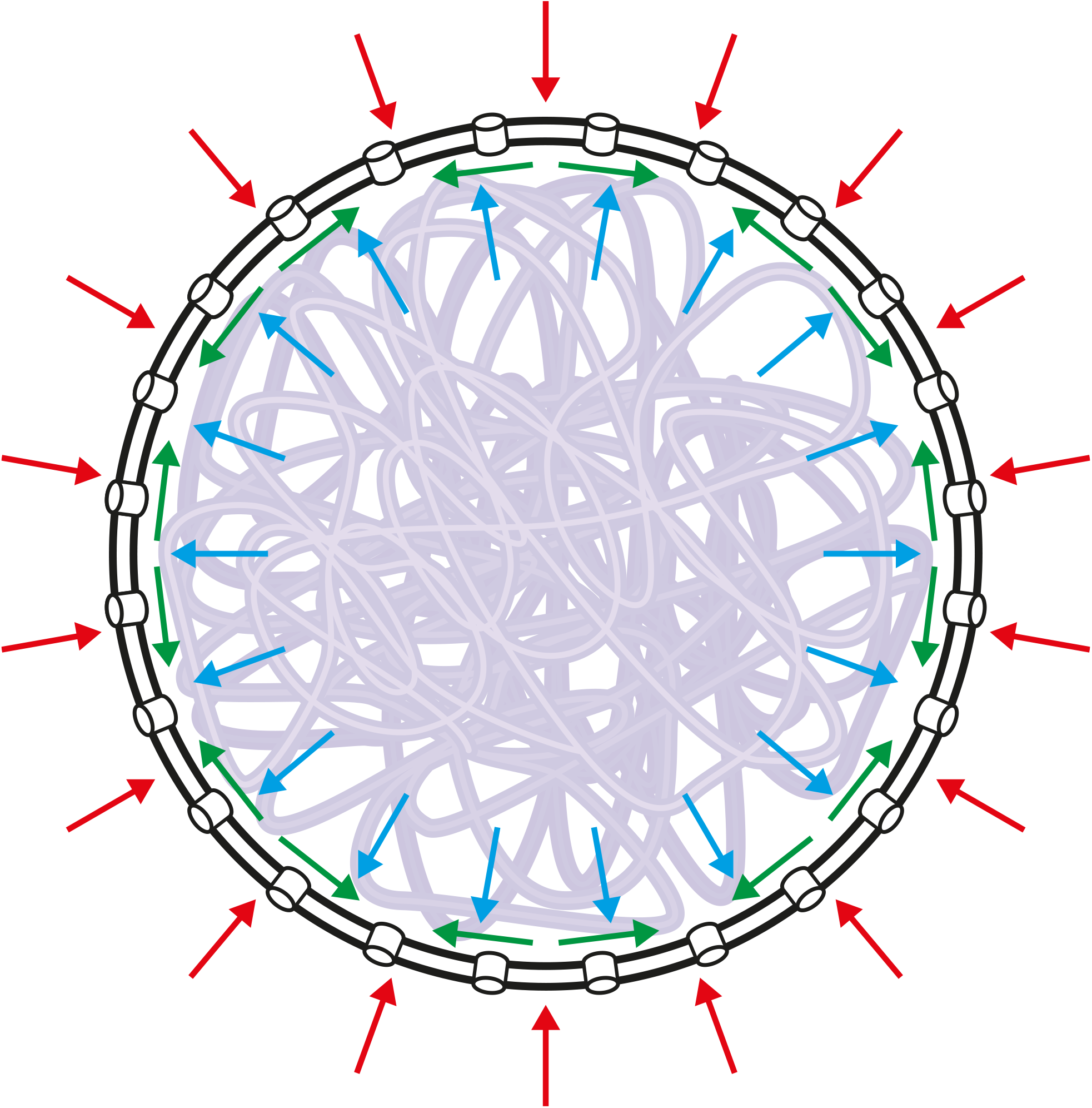
Force balance diagram of a spherical nucleus: The nuclear envelope (NE), depicted by two concentric black circles (representing the inner (bilayer that in most species is attached to a lamin layer) and outer NE membranes) perforated by channels (representing the nuclear pore complexes), envelopes the chromatin (semitransparent purple curve). The red arrows represent the sum of the inward pressures that tend to compress the nucleus and the blue arrows represent the sum of the outward pressures that tend to expand the nucleus. In the case where the outward pressures are larger than the inward pressures, the difference of the pressure is balanced by NE tension, depicted by green arrows whose resultant (vector sum) gives rise to an inward force, which limits the expansion of the nucleus.

**Table 1:** Order of magnitude estimates of the contributions of different nuclear and cytoplasmic components to the inward and outward pressures that are exerted on the nucleus.

| Species/Source of pressure | Chromatin polymer | Chromatin counterions | Cytoskeletal elasticity | Cytoplasmic proteins | Nucleoplasmic proteins |
| --- | --- | --- | --- | --- | --- |
| Human | 30 Pa | 300 Pa | 1-100 Pa | 8000 Pa | 8000 Pa |
| Fission yeast | 1.5 Pa | 20 Pa | 1-100 Pa | 8000 Pa | 8000 Pa |

### 2.1 Polymer model of the osmotic pressure of the chromatin

Polymer physics predicts that the chromatin macromolecule has a thermodynamically preferred radius which it would occupy if not constrained in any way; this is known as the radius of gyration and denoted by *R_g_*. If the radius of the nucleus *R* is smaller than *R_g_*, then the chromatin is confined by the NE to a volume that is smaller than its preferred volume, which results in an outward osmotic pressure *p_p_* exerted by the chromatin on the inner side of the NE. This contribution to the outward pressure increases with the size of the genome, if the size of the nucleus remains the same. The radius of gyration of a chromatin macromolecule is determined by the length of the DNA molecule that constitutes it, its degree of compaction in the chromatin fiber, and the persistence length of the chromatin fiber. In addition to these, self-attraction or repulsion (including steric, excluded volume) of the chromatin fiber result in different dependence of *R_g_* on the genome size. For example, self-attraction can lead to chromatin collapse (as in [20, 21]). Since we are interested in identifying the dominant forces, we estimate an upper bound on the possible values of *R_g_* and subsequently the pressures *p_p_*. This occurs for the case where the chromatin fiber is in a “good solvent”, namely with chromatin-chromatin steric repulsion that is larger than its self-attraction, so that the radius of gyration is then maximal [22].

In that situation, the radius of gyration of a chromatin macromolecule is *R_g_* ≈ *ℓ_p_*(*L/ℓ_p_*)^3/5^ [22], where *ℓ_p_* is the persistence length of the chromatin, and *L* is the contour length of the chromatin, which is the length of the DNA macromolecule within the chromatin divided by the chromatin linear density. Both the density of the chromatin fiber and its persistence length depend on the structure of the chromatin fiber which is generally not uniform [23]. However, different experimental measurements and theoretical estimates of the chromatin density and persistence length result in a range of 7 – 11nm/kb for the chromatin density and *ℓ_p_* ~ 30 – 180 nm for its persistence length [24, 25]. Because we are interested in calculating an upper bound for the pressure, we choose the parameters that yield the maximal radius of gyration: a persistence length of 180 nm and a chromatin density of 11 nm/kb. Furthermore, we treat the entire genome as one chromosome (chromatin macromolecules) instead as made up of multiple chromosomes, which simplifies the calculation and results in a slightly increased pressure (in the spirit of finding an upper bound); this is because the preferred volume of a chromosome, *V_p_*, scales as 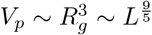, so that the preferred volume of two concatenated chromatin molecules is larger than the sum of the preferred volumes of the two separate chromatin molecules, since 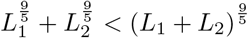.

When the entire genome is treated as one chromatin macromolecule, the contour length of the chromatin molecule is calculated directly from the genome size and the linear density of the chromatin. The diploid S. pombe genome comprises 27.6 Mbp of DNA [26], which gives rise to a contour length of approximately 300 μm, while the diploid human genome comprises 6.4 Gbp [27], which gives rise to a contour length of approximately 70 mm. Substitution of the contour lengths of human and yeast cells and the persistence length of the chromatin, yields chromatin radii of gyration of approximately 400 μm and 15 μm, respectively. Both of these values are significantly larger than their respective nuclear radii of 6.2 μm and 1.6 μm, calculated from respective nuclear volumes of *V* ≈ 1000 μm^3^ [28] and *V* ≈ 17 μm^3^ [8] for idealized spherical nuclei. Therefore, in both cases the confinement of the chromatin (assumed to be in a good solvent for the calculation of an upper bound) in the nucleus will contribute an outward pressure. For such a polymer, with radius of gyration *R_g_* confined in a sphere of radius *R* < *R_g_*, the compression energy Δ*F* is given from polymer physics theory by Δ*F* = *k_B_T* (*R_g_/R*)^15/4^, where *k_B_T* is the thermal energy [29]. The pressure *p_p_* is then given by the negative derivative of the compression energy Δ*F* with respect to the volume of the nucleus, *V* = 4*πR*^3^/3, which results in the following expression

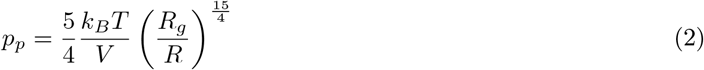

Substitution of the values of *V*, *R*, and *R_g_* for the two species results in an order of magnitude estimation for the outward pressure due to confinement of the chromatin polymer of ≈ 30 Pa for human nuclei and ≈ 1.5 Pa for S. pombe cells. As we show below, these upper-bound values are still far below the pressures due to preferentially-localized proteins (see Table 1).

We note that we treated the chromatin as a neutral polymer while in fact, it is negatively charged. This treatment is justified because the screening length (the decay length of the electrostatic interactions in electrolyte solution) in cellular systems is ≈ 0.7 nm [30], which is very small compared with the polymer radius of gyration and the nuclear size. However, the net electric charge of the chromatin can contribute to the outward pressure indirectly, due to the localization of counterions in the vicinity of the chromatin, as we now discuss.

### 2.2 Osmotic pressure of the chromatin counterions

Chromatin is a complex of a negatively charged DNA polymer that links and is wrapped around positively charged histone proteins. Due to the steric and polymeric constraints involved, chromatin is not neutral has a net negative charge. As a result, the chromatin macromolecule is surrounded by a “cloud” of soluble, positive counterions that neutralize its charge by being localized within the nucleus despite being able to diffuse through the NPC [31]. Theoretical studies of analogous solutions of charged colloids [32] suggest that depending on the detailed molecular structure and charge of the chromatin and the concentration of electrolytes in the nucleoplasm, the cloud of counterions may be either highly localized near the chromatin fiber, or dispersed within the nucleoplasm. Localized counterions are constrained to move together with the chromatin and thus present a negligible contribution to the osmotic pressure. On the other hand, the contribution of dispersed counterions can be significant, similar, in the limiting case, to that of an ideal gas, in which all the counterions can collide with the NE.

Since we are interested in an upper bound, order of magnitude estimate of the contribution of the counterions to the osmotic pressure, we consider the limiting case in which the counterions are dispersed within the nucleoplasm and contribute to the pressure as an ideal gas. In this limit, the concentration of counterions is uniform within the nucleoplasm and is determined by two conditions. The first is equality of electrochemical potentials of each of the different species of counterions. The ions diffuse from the nucleus to the cytoplasm (and vice versa) via the nuclear pore complexes which allows them to reach chemical equilibrium; this would not be the case if the NE contained active, ion pumps that directly transport ions between the cytoplasm and nucleoplasm[33]. The second condition is that the cytoplasm and nucleoplasm are each to a very good approximation electro-neutral, which arises from minimization of electrostatic energy [31]. In a simple model that includes one type of monovalent cation, one type of monovalent anion, and a negatively charged polyelectrolyte (chromatin), these two condition give rise to the following expression for the outward pressure of the chromatin counterions *p_c_* (see Appendix I):

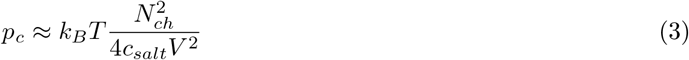

where *k_B_T* is the thermal energy, *N_ch_* is the overall charge of the chromatin in units of electron charges, *c_salt_* is the intracellular salt concentration, and *V* is the nuclear volume. We note that we do not include the charges of other proteins in the nucleoplasm and cytoplasm since we expect them to be negligible compared to the highly charged chromatin. Furthermore, since we are seeking an upper bound on this quantity, we do not include the effect of the negatively charged cytoskeleton [34], which cancels some of the pressure difference, leading to a net, outward pressure that is even smaller than that given by Eq. 3.

To calculate the net charge of the chromatin, *N_ch_*, we calculate the number of histone proteins based on the genome size (6.4 Gbp in humans cells and 27.6 Mbp in S. pombe cells [26, 27]) and a typical nucleosome density of one per 200 bp [35]. We then subtract the total positive charge of all the histones, which is the product of the number of histones and the net positive charge of the histone octamer proteins, 220 electron charges [35], from the negative charge of the DNA (based on a negative charge of 2 electron charges per base pair) for both the linker DNA and the nucleosomal DNA. This results in a total chromatin charge of *N_ch_* ≈ 5.8 · 10^9^ electron charges for humans and *N_ch_* ≈ 2.5 · 10^7^ for S. pombe. We substitute these values into Eq. 3, along with a typical intracellular salt concentrations of ≈ 200 mM [27, 36] and nuclear volumes (~1000 μm^3^ [28] for human nuclei and ~17 μm^3^ for S. pombe nuclei [8]), which results in an approximate upper bound for the *p_c_* of the two species: 300 Pa for humans and 20 Pa for yeasts. As shown in Table 1, these values are much smaller than those associated with the localized proteins.

### 2.3 Cytoskeletal elastic pressure

The nucleus is tethered to the cytoskeleton by the LINC complex [37], which is embedded in the NE. This couples mechanical forces in the cytoskeleton and the LINC complexes, thereby exerting elastic forces on the NE, which may contribute to either outward or inward pressures. To estimate the magnitude of the cytoskeletal pressure on the nucleus, we expect this pressure to be of the same order of magnitude as elastic stresses within the cytoskeleton that are mechanically balanced by its own rigidity response. Atomic force microscopy measurements show that the stiffness of the cytoskeleton is of the order of 1-100 pascals [28, 38], which suggest the cytoskeletal pressure is of the same order. We note that depending on the structure of the cytoskeleton and its anchoring to the extra-cellular matrix the cytoskeletal forces may be inward (e.g. compression of the nucleus that contacts the actin cortex in spread cells) or outward (e.g., stretch of the nucleus by sarcomeres in muscle cells or stress fibers in other cells). However, the sign of the pressure does not play a role in determination of the nucleus volume. This is because, as we show in the subsection below, the estimated magnitude of this contribution to the inward or outward pressures is negligible compared to the contributions of the osmotic pressure of proteins localized to the cytoplasm and those localized to the nucleoplasm due to active, nucleocytoplasmic transport.

### 2.4 Osmotic pressures of proteins preferentially-localized to the cytoplasm and nucleoplasm

The average concentration of cellular proteins in human and yeast cells is about 2.6 · 10^6^ proteins/μm^3^ [39], which in a homogeneous, ideal solution, contributes ≈ 10 kPa to the osmotic pressure of the solution. Since about 80% of the proteins are preferentially localized, the results of the SI Appendix “Localized protein pressure” indicates that the osmotic pressures of the localized proteins in the cytoplasm and nucleoplasm is of the order of 8 kPa. However, as we now explain, not all of the these ≈ 10 kPa of osmotic pressure are contributed to the inward or outward pressure of the respective compartment of the proteins.

The distribution in the two compartments of proteins that can freely cross the NE by passive diffusion through the NPCs is determined by equilibrium thermodynamics. In the ideal solution limit, this dictates that the concentrations of any protein species across the NE are equal, so that their contributions to the osmotic pressures of the two compartments are equal. Therefore, no imbalance of the inward or outward pressure can ever be form that lead to changes of the compartments volume. An active, non-equilibrium process is thus required to partially localized proteins to either compartment and prevent their free exchange, so that *these* proteins contribute to the net osmotic pressure imbalance that drives fluid flow which effectively determine nuclear volume.

Nucleocytoplasmic transport is an active, out-of-equilibrium process that preferentially localizes proteins to either the cytoplasmic or nucleoplasmic compartment (e.g. related to non-equilibrium Ran-GTP gradients [40, 41, 42]). Localization of proteins to a given compartment means that the steady-state concentration of those proteins in that compartment is larger than their concentration in the other. This can be seen from the condition (for equal fluxes in steady state) that the rate of protein transport into the nucleus must equal the rate of protein transport out of the nucleus. In the spirit of first order kinetics, the equal flux condition is quantified as *k_i_c_c_* = *k_e_c_n_*, where *k_i_* and *k_e_* are the nuclear import and export rates of a given protein species, respectively, and *c_c_* and *c_n_* are its respective cytoplasmic and nucleoplasmic concentrations. Under conditions of chemical equilibrium, the absence of active protein transport implies the two rates are equal, *k_i_* = *k_e_*, since both are determined by diffusion through the nuclear pores; this implies that *c_c_* = *c_n_*. However, active transport can result in unequal import and export rates, which results in unequal concentrations in the nucleus and cytoplasm, *c_c_* ≠ *c_n_* – i.e., preferential localization of certain proteins to one of those compartments. This argument is further quantified in the SI section “Localized protein pressure”. The osmotic pressures of the preferentially-localized proteins in the nucleus and cytoplasm are not equal, in contrast to equilibrium. As we show, those contributions are dominant (see Table 1) and the difference in osmotic pressures must, in general, be balanced by the surface tension of the NE as in Eq. 1.

However, in many cases, the NE is relaxed (see the next subsection and the Discussion) with effectively zero tension. Mechanical equilibrium then dictates that, the sum of the contributions of all the localized proteins to the inward and outward pressures must be equal (see the section on “Model and results” section below). Deviations from the equality of inward and outward pressures, such as transiently caused by production of new proteins, induce fluid flow that changes the nuclear volume and restores the equality of pressures. This is to be contrasted with the equilibrium case where proteins passively diffuse through the NPC and reach equal concentrations, in which their inward and outward pressures are equalized by flow of proteins rather than fluid. Thus, in the equilibrium case, nuclear volume changes are not induced.

An estimate of the fraction of preferentially-localized proteins (due to non-equilibrium transport) relative to the total number of proteins is given by a Study of X. laevis oocytes show that 83% of cellular proteins are preferentially-localized to either the cytoplasm or the nucleoplasm [43]. This was determined by physical separation of the nucleus and cytoplasm which is possible due to the large nucleus size of the X. laevis oocytes. Determining this for other cell types with smaller nuclei is difficult because most nuclear fractionation protocols require the use of detergents that damage the NE and mix the nucleoplasm and cytoplasm. However, since the localization of proteins is usually associated with their function, one can expect that the homologs of the X. laevis proteins in yeast and human cells are similarly localized. We thus expect that the percentage of localized proteins in yeast and human cells is also of the order of 80%. In the non-equilibrium situation, in the special case that the NE is relaxed, the inward/outward pressures are equal to the osmotic pressure of a protein solution, whose concentration is the average, cellular concentration of all localized proteins (see SI Appendix “Localized protein pressure”). Therefore, we estimate that 80% of the aforementioned 10 kPa of the overall osmotic pressure of all proteins is contributed to the inward and outward pressure of the cytoplasm and nucleoplasm, namely that the upper bound for the contribution is 8 kPa.This allows us to estimate the contribution of the localized proteins to the \osmotic pressures of the cytoplasm and nucleoplasm in humans and yeast cells: The average concentration of cellular proteins in human and yeast cells is about 2.6 · 10^6^ proteins/μm^3^ [39], which in a homogeneous, ideal solution, contributes ≈ 10 kPa to the osmotic pressure of the solution. Since about 80% of the proteins are preferentially localized, the results of the SI Appendix “Localized protein pressure” indicates that the osmotic pressures of the localized proteins in the cytoplasm and nucleoplasm is of the order of 8 kPa.

It is important to note that the actual contribution of the localized proteins to the inward and outward pressures is less than the estimate of 8 kPa due to a number of factors. First, preferential localization, rather than complete localization, results in a lower net contribution of the protein species to the inward and outward pressure (see SI section “Localized protein pressure”). Second, the estimate of 80% localized proteins refers to the percentage of the protein species rather than to the number of proteins. Furthermore, this quantity was measured in the special cell-type of oocytes rather than somatic cells. Therefore, the actual contribution in somatic cells may be smaller. Third, attractive interactions between proteins, such as those that lead to oligomerization of the proteins, reduces their contribution to the osmotic pressure in a way that is not captured by the ideal solution approximation for the osmotic pressure. Therefore, our estimate of an 8 kPa contribution of the localized proteins to the inward and outward pressures should be considered as an upper limit. Nonetheless, even if this estimate is reduced by an order of magnitude to 0.8 kPa, it still much larger than all the other contributions that appear in Table. 1. This strongly suggests that the pressures due to localized proteins is the major factor that determines nuclear size. We now use these ideas to formulate a quantitative model for the volumes of the cell and the nucleus in the section “Model and results”, where we also discuss their proportionality. Before that, we briefly discuss NE tension in terms of its the non-linear mechanical response to expansion of its area.

### 2.5 NE tension

The Young-Laplace law expresses how the tension of the NE (resultant of the vector forces in the plane of the membrane, see Fig. 1) results in a net inward force that in mechanical equilibrium, balances the net outward pressure of the nucleoplasm. The NE comprises two lipid membranes and, in most organisms, at meshwork of semi-flexible filamentous proteins (lamina) [44, 45]. Since the mechanical response of both the lipid membranes and the semi-flexible polymers to stretching is highly non-linear and stiffens with increasing expansive strain [46, 47], these properties are expected to be reflected in the mechanical response of the NE: When the nuclear volume is relatively small (as can be the case in a hypertonic environment), the NE is relaxed and undulated, expansion only reduces the entropy of the undulations and is typically small [48, 49]. In contrast, when the nuclear volume is relatively large (hypotonic environment), the NE is taut. This expansion of the membrane changes the conformations of its constituents (e.g., packing density of lipids or lamin fiber structure) and thus raises the membrane energy and increasing its tension. Therefore, following the theoretical description in [50], we approximate the NE tension *σ* in its relaxed state by *σ* = 0 and in its taut state, (when the nuclear radius exceeds a characteristic radius *R* > *R_c_*) by a positive value *σ* which itself depends on the amount of stretching and is thus a function of the nuclear radius *R*. The characteristic radius *R_c_* is determined by the number of molecules (e.g., lipids, lamin proteins) that comprise the NE and the molecular details of the NE structure. The detailed dependence of *R_c_* on the number of molecules, as well as the dependence of *σ* on the nuclear radius *R* for *R* > *R_c_*, involve many molecular details of the NE, which are outside of the scope of this paper. However, the relaxed and taut states can easily be distinguished experimentally, since the relaxed NE is undulated and is therefore less spherical than the taut NE, which is smooth.

## 3 Model and results

Table 1 summarizes the order of magnitude estimates of the different inward and outward pressures discussed above and suggests that the dominant contributions are the osmotic pressures due to localized proteins in the cytoplasm and nucleoplasm. These osmotic pressures are estimated to be about an order of magnitude larger than inward or outward pressures from the other sources, which themselves were upper bounds. This motivates us to consider a minimal model for determination of the volumes of the cell and the nucleus that includes the osmotic pressures due to soluble factors but neglects the effects of chromatin and cytoskeleton. While the chromatin and cytoskeleton may help control the cellular and nuclear shapes, their relatively small pressures only perturb the volumes of the cytoplasm and nucleoplasm determined by mechanical balance of osmotic pressure of the soluble factors.

Our steady-state model considers two nested compartments (cytoplasm and nucleus) that contain both localized (large proteins and RNA that are actively transported, as discussed above) and non-localized solutes (small molecules/proteins and ions that freely diffuse through the nuclear pores) (see Fig. 2). During steady state (e.g. interphase in non-dividing cells), the volumes of the compartments are constant in time, which implies that that they are in mechanical equilibrium, but not necessarily thermodynamic equilibrium. The outer (plasma) membrane separates the cytoplasm from the extra-cellular environment and allows, at sufficiently long timescales, free flow of water and diffusion of ions via channels, but prevents free diffusion of large solutes. In addition, the plasma membrane also contains mechanosensitive pumps that regulate the transport of small molecules and ions so that to a first approximation, the tension of this membrane can be neglected [33]. In contrast to the outer membrane, the inner membrane (NE) which separates between the cytoplasm and nucleoplasm is a semi-permeable membrane that allows free flow of water and diffusion of small solutes (e.g., small molecules/proteins and ions), without any active transport via pumps. As explained above, larger solutes (e.g., proteins, RNA) are actively transported through the nuclear pore complexes (NPCs) and thus are preferentially-localized to one of the compartments.

**Figure 2:**
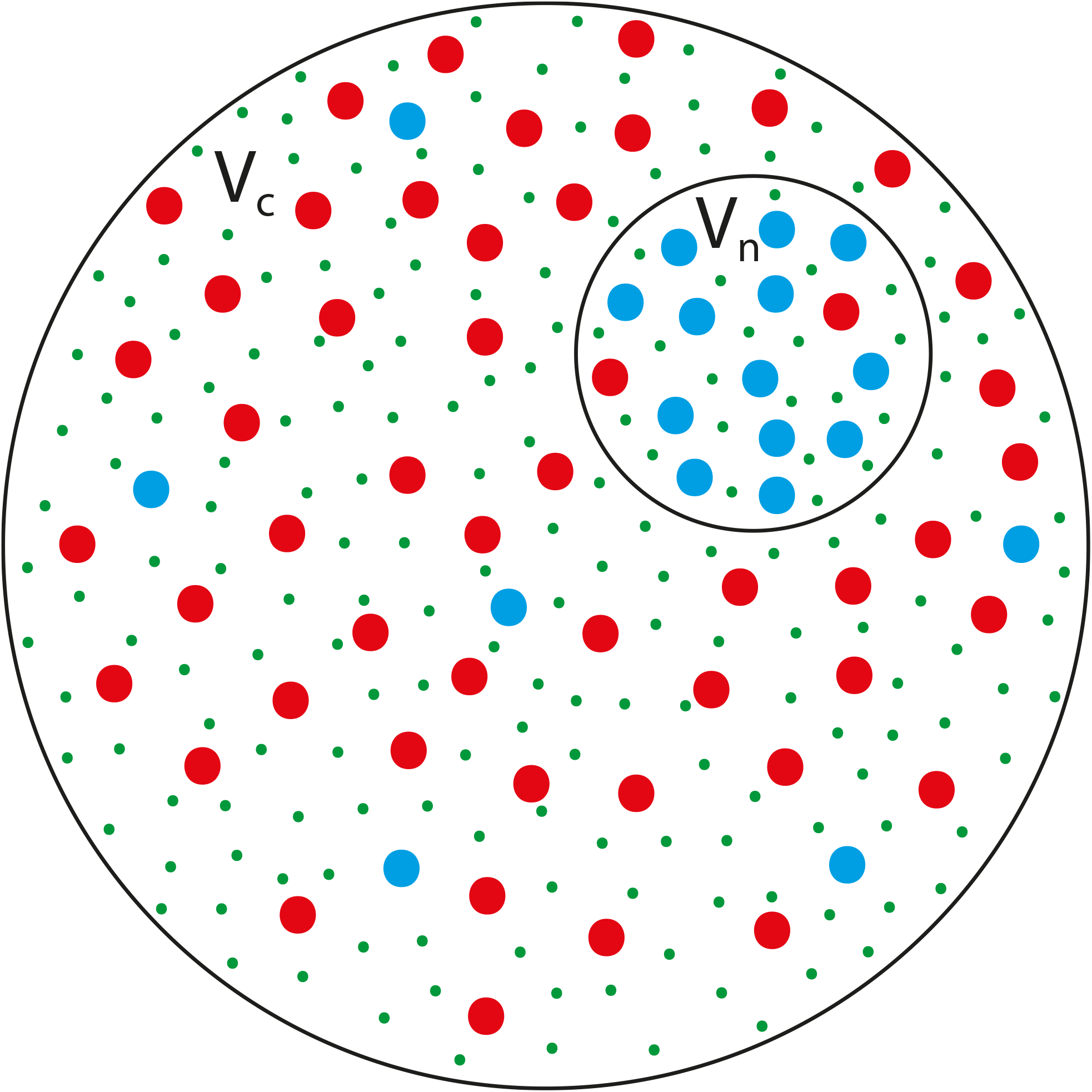
A schematic cartoon of the proposed model: Two black circles representing membranes that delineate two compartments, the cytoplasm, whose volume is *V_c_* and the nucleoplasm whose volume is *V_n_*. The outer circle represents the plasma membrane which separates the cytoplasm from the extra-cellular environment and does not allow free diffusion of large solutes (red, blue); steady-state exchange of small molecules and ions with the cellular environment via active pumps regulates the concentration of those small solutes or ions that may also passively diffuse through channels (green dots). The inner circle represents the nuclear envelope, a semipermeable double membrane (that in some species is attached to the lamina) that separates the cytoplasm and the nucleoplasm and allows free diffusion of small solutes (small green dots) but not of large solutes (large red and blue dots) through the nuclear pores. Due the free diffusion of the small solutes, their concentration in the two compartments is the same. In contrast, the large solutes which are transported across the NE by active mechanisms are not equally distributed in the nucleus and cytoplasm. The species of large solutes represented by the blue dots are preferentially-localized to the nucleoplasm while the species of large solutes that are represented by the red dots are preferentially-localized to the cytoplasm. This situation is treated in the SI appendix “Relation to nucleocytoplasmic transport”, and in the main text we treat a minimal model where the localization is complete; as we show in the SI, the two cases yield similar results.

In our model, we account for the nucleocytoplasmic transport by considering the simple situation of a single species of 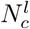 solute molecules that are “confined” to the cytoplasm and 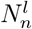 solute molecules confined to the nucleoplasm. Neither species can cross the nuclear pore complexes (NPCs). These solutes are completely localized to their respective compartments, and their number is thus conserved with each compartment. This approach is justified in the SI Appendix “Relation to nucleocytoplasmic transport”, where we show that it yields the same results as the realistic case where a multitude of protein species are only preferentially-localized rather than completely localized; for clarity, we present in the main text this minimal model which is easier to solve.

Mechanical equilibrium across a membrane is described by Eq. 1. In our model, the pressures on the left hand side of Eq. 1 are the osmotic pressures of the various solutes. Since the small, non-localized solutes can freely diffuse across the NE, their osmotic pressures on the two sides are equal and cancel each other on the left hand side of Eq. 1. Thus, only the contributions of the completely-localized, large solutes remain, which depends on 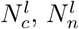, and the free volumes of the cytoplasm and the nucleoplasm:

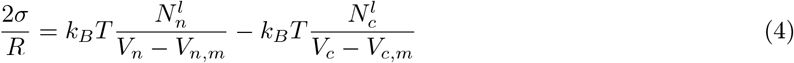

where *σ* is the NE tension, *R* is the radius of the spherical nucleus, *k_B_T* is the thermal energy, *V_n_* and *V_c_* are the respective volumes of the nucleoplasm and cytoplasm, and *V_n,m_* and *V_c,m_* are their respective minimal volumes (the total volume of all non-solvent molecules in the compartment). The free volumes, *V_c_* – *V_c,m_* and *V_n_* – *V_n,m_* are used here and in the rest of the paper to calculate concentrations in order to account for the steric, excluded volume interactions among the solutes, whose overall volume may be significant [28].

While the non-localized, small solutes (small molecules and ions) that freely diffuse across the NE do not play a role in determining mechanical force balance across the NE, this is not the case for the plasma membrane. The mechanical balance across the plasma membrane involves the difference of the osmotic pressure of the extra-cellular environment and the total osmotic pressure in the cytoplasm, which has contributions for both localized and non-localized solutes. To find the total osmotic pressure in the cytoplasm, we must first determine the total number of solutes within it. The small, diffusive, non-localized solutes in the cell, are found in both the cytoplasm and nucleoplasm. We denote the numbers of non-localized solutes found in each respectively by by *N_c_* and *N_n_*, and define their total number: *N* = *N_c_* + *N_n_*. Since these small solutes can freely cross the NE, they are in chemical equilibrium and their freely-diffusing concentrations in the cytoplasm and nucleoplasm are equal, as are their contributions to the osmotic pressures of the cytoplasm and nucleoplasm, so that *N_c_*/ (*V_c_* – *V_c,m_*) = *N_n_*/ (*V_n_* – *V_n,m_*). This equation and the equation for the total number of non-localized, small solutes, *N* = *N_c_* + *N_n_*, can be solved to give the number of non-localized, small solutes in each compartment:

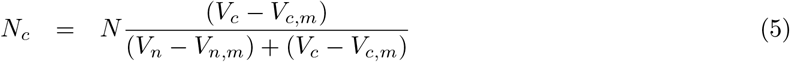

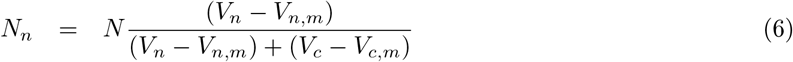

Henceforth, for brevity, we refer to these as small solutes and omit the word non-localized.

The plasma membrane includes mechanosensitive ion pumps that regulate the total number of small, cellular solutes, *N*, such that to a first approximation, the tension on the plasma membrane is zero (*σ* = 0) [33]. Therefore, when applied to the plasma membrane, Eq. 1 suggests that the osmotic pressure of the extra-cellular environment, which is determined by external conditions, and the osmotic pressure of the cytoplasm, due to both the non-localized and completely-localized solutes, *N_c_* and 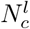, are equal, to a first approximation. Using Eq. 5 for *N_c_*, we can explicitly write the osmotic pressure in the cytoplasm, which is 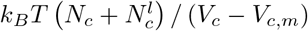, in terms of the total number of cytoplasmic non-localized and completelylized solutes, so that Eq. 1 for the plasma membrane becomes:

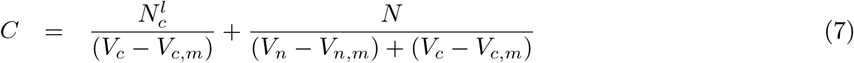

where *C* is the solute concentration in the extra-cellular environment.

Eqs. 4 and 7 relate the volume of the cytoplasm and that of the nucleoplasm to the NE tension *σ*. Since there are two equations and three variables (two volumes and the tension), this system of equations cannot be solved uniquely to determine the volumes of the cytoplasm and nucleoplasm without additional equation that relates the tension to the two volumes, *σ*(*V_n_, V_c_*). As explained in the subsection “NE tension” above, *σ* is a non-linear function of the nuclear size, whose exact form depends on molecular details of the NE. However, as explained in the subsection “NE tension”, due to the non-linear, elastic properties of the NE, when the area of the NE (hence the nuclear radius) is smaller than a characteristic value, the NE is relaxed and its tension is negligible. In this important case, which we consider in the subsection below, Eqs. 4 and 7 can be solved for *V_c_* and *V_n_*. The case where the NE is taut (its area is larger than the characteristic value) is considered in the Discussion section below.

### 3.1 Cytoplasmic and nuclear volumes are proportional for relaxed NE (*σ* = 0) in the ideal solution limit

For *σ* = 0, Eq. 4 indicates that the ratio between the free volumes of the cytoplasm and nucleoplasm equals to the ratio of the numbers of completely-localized solutes in the two compartments

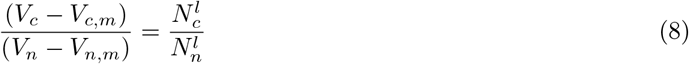

In the “ideal solution” limit where the minimal (steric, excluded) volumes *V_c,m_* and *V_n,m_* are negligible compared with the cytoplasmic and nuclear volumes respectively, Eq. 8 predicts that the ratio of the nuclear and cell volumes are simply proportional to the ratio of the numbers of completely-localized, large solutes in each. This is thus independent of genome size, cytoskeletal properties, and electrostatics of the cell, consistent with the observation that unless the nuclear transport or protein expression is changed, the volume ratios of the nucleus and cytoplasm are constants. Of course, this “ideal solution” is only a gross approximation since the minimal volumes of the cytoplasm and nucleoplasm may be of the same order of magnitude as the entire volumes of each of the compartments [28]. However, in Appendix “Cytoplasmic and nuclear volumes in the non-ideal limit” of the SI, we show that even if the minimal volumes are included, the ratio of the two volumes is still a constant, in the approximation that the volumes of the non-diffusive structures (e.g. cytoskeleton and chromatin) are negligible compared to the total volume of their respective compartments. As our estimates presented in Appendix “Cytoplasmic and nuclear volumes in the non-ideal limit” show, this is indeed the common biological case. Therefore, we henceforth focus on the case where *V_c,m_* ≪ *V_c_* and *V_n,m_* ≪ *V_n_* so that *V_c_* – *V_c,m_* and *V_n_* – *V_n,m_* are well approximated by *V_c_* and *V_n_*, respectively. The detailed results for the case where *V_c,m_* and *V_n,m_* are included are presented in Appendix “Cytoplasmic and nuclear volumes in the non-ideal limit” of the SI and yield the same qualitative conclusion.

In the ideal solution limit, Eq. 8, together with Eq. 7 for the mechanical balance of the plasma membrane determine the volumes of the cytoplasm and nucleoplasm for the case where *σ* = 0

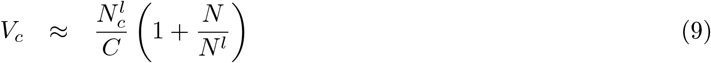

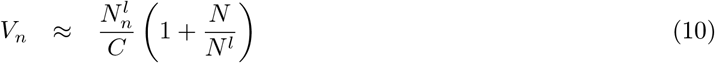

where *N* is the total number of non-localized, small solutes, and 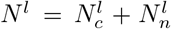 is the total number of completely-localized, large solutes (in both the cytoplasm and nucleoplasm).

## 4 Discussion

The quantitative estimates of the various contributions to the osmotic pressures of the nucleus and the physics-based model that predicts the proportionality of the nuclear and cellular volumes for many common cases, is a physical and quantitative parallel of the qualitative, biology-based discussion in [6]. Our estimates of the osmotic pressures includes the contributions of diffusive solutes and non-diffusive structures such as the chromatin and cytoskeleton. We show that the factors that determine the mechanical balance across the NE are the osmotic pressures of proteins preferentially-localized to either the cytoplasm or the nucleus (due to active transport). Thus, our minimal model neglects the pressures that originate in are the chromatin or cytoskeleton, which are estimated (see Table 1) to be about an order of magnitude smaller, at least, than the osmotic pressures of the solutes.

The force balances we consider, 4 and 7, have a simple solution for the case that the NE is relaxed, for which the volumes of the cytoplasm and nucleoplasm are respectively given by Eqs. 9 and 10. Our model predicts that the ratio of the volume of the nucleoplasm, *V_n_*, and the volume of the cytoplasm, *V_c_*, is proportional to the respective ratios of the number of localized proteins in each compartment: 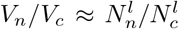. This prediction suggests that as long as the NE is relaxed, the nucleoplasm-to-cytoplasm volume ratio (NC ratio) is governed almost exclusively by the active, nucleocytoplasmic transport of proteins and RNA molecules to and from the cytoplasm and nucleoplasm. Biophysical or biochemical perturbations of the cell or its environment that do not change nucleocytoplasmic transport, such as osmolarity of the cellular environment or variations in the amount of chromatin, will not change the NC ratio as long as the NE remains relaxed, in agreement with multiple experiments [4, 8, 9, 28, 50]. In contrast, we predict that genetic or chemical perturbations of components implicated in nucleocytoplasmic transport will change the NC ratio, in agreement with recent findings [6, 8, 51, 52, 53, 54, 55]. The implication of nucleocytoplasmic transport as the main governing factor of the NC ratio may also explain how indirect biological effects reviewed in [6], such as cytoplasmic factors, transcription and RNA processing, and the LINC complex, can affect nuclear size. Cytoplasmic factors, transcription and RNA processing are responsible for the presence of the large solutes that are localized by nucleocytoplasmic transport, while some of the components of the LINC complex are implicated in the nucleocytoplasmic transport itself [56].

Furthermore, even when the nucleus is not spherical due to the action of cytoskeletal shear forces or external mechanical forces, our theory predicts that for a relaxed NE, the NC ratio is constant. As explained at the beginning of the section “Physical forces that influence nuclear volume”, in non-spherical nuclei the right hand side of the force balance Eq. 1 is changed so that the local curvature of the NE replaces the inverse of the nuclear radius. However, when *σ* = 0, the shape of the nucleus does not enter the force balance equation and does not affect the predicted volumes. In that case, we predict that the shear forces determine nuclear shape, while the pressure differences determine nuclear volume. This prediction for non-spherical nuclei agrees with the experimental observation that the NC ratio was maintained in adhered cells despite changes of the substrate area [28], and stiffness [57], which modified both the nuclear and cellular shapes and volumes.

In fact, the robustness of the NC ratio to various biophysical and biochemical perturbations is an observation that is more than a century old [7]. It has been observed in wide variety of organisms, but was not explained mechanistically [6] as we have now proposed. The ubiquity of the constancy of the NC ratio suggests that in most physiological scenarios, the NE is relaxed, a property that may be favored because it reduces nuclear rupture [58], and subsequent DNA damage [19]. However, in non-physiological, laboratory conditions, our theory predicts that it is possible for the NE to become taut – for example, under hypotonic conditions. Eq. 10, which predicts the nuclear volume for the case of relaxed NE, suggests that the nuclear volume increases with decreasing osmolarity of the extra-cellular environment. As the volume of the nucleus increases, the surface area of the NE must increase as well, eventually stretching the NE and rendering it taut. In that case, the NE tension becomes important, and for spherical nuclei, the volumes of the cytoplasm and nucleoplasm, and the NE tension, are related by the Eqs. 4 and 7, together with the expressions for the minimal volumes *V_c,m_* and *V_n,m_* given in Appendix “Cytoplasmic and nuclear volumes in the non-ideal limit” of the SI. If the relation between the nuclear radius and the NE tension *σ*(*R*) is known or experimentally measured, these equations can be solved to predict the volumes of the cytoplasm and nucleoplasm as a function of the extra-cellular osmolarity *C*. The NC ratio is then not constant. For non-spherical nuclei, such as in adhered cells, the equations are more complex since, as explained above, the mechanical balance that determines nuclear volumes depends on the nuclear shape and is thus no longer independent of the shear forces. In either case, the NC ratio is expected to depend on the extracellular osmolarity *C* rather than being constant, as indeed observed in cells in hypotonic conditions [50].

Although predicting the dependence of the NE tension on the nuclear radius, *σ*(*R*), is outside of the scope of our work, our discussion in the section on “NE tension” provides a qualitative prediction regarding the transition between the relaxed and taut regimes of the NE. As explained there, the mechanical properties of the NE are highly non-linear: In the relaxed state, stretching the NE mainly suppresses undulations while in the taut regime, stretching the NE increases the separation between the molecules that constitute the NE. The transition between these two regimes occurs when the NE first becomes smooth, a state in which the area of the NE is proportional to the amount of NE components, which are in their native, unstretched molecular conformation. This implies that the NE area at which this transitions happens scales with the amount of the NE components. We therefore expect that inhibition of the production of NE components may promote a transition of the NE to a taut state, thereby limiting nuclear growth and changing the NC ratio. Indeed, inhibition of nuclear growth due to lack of NE components was observed in mutant fission yeast cells whose nuclear export of RNA and lipid production were both inhibited [52], and in an extract of X. laevis nuclei that expanded in response to titration (addition) of lamin proteins [59]. The fact that both of these experiments represent non-physiological conditions (defective nuclear export and absence of cytoplasm), also supports the idea that in most physiological conditions the NE is relaxed.

Our theory can be directly tested by measurements [60] of the total number of large solutes that are localized to the cytoplasm and nucleoplasm, 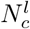 and 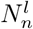, and comparing the ratio of the cytoplasm and nucleoplasm volumes to the ratio of 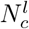 and 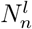. However, accurate measurements of 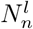 and 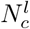 are experimentally challenging because nuclear fractionation protocols commonly use detergents that perforate the NE membranes thereby mixing the nucleoplasm and the cytoplasm [43]. In X. laevis oocytes, the large sizes of the nucleus and the cell allow the nucleus to be removed mechanically without perturbing the NE membranes [43]. This potentially may allow measurements of the nuclear and cytoplasmic protein concentrations using mass spectroscopy and calculation of the values of 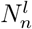 and 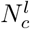 (see SI Appendix “Relation to nucleocytoplasmic transport”), thus providing a quantitative test of our predictions. Furthermore, qualitative tests of our theory can be conducted by changing the localization of abundant, large soluble proteins by fusing them with nuclear localization signals or nuclear export signals; our theory predicts that in this case the NC ratio will then increase or decrease, respectively. In addition, cellular events that change the proteome and its localization, such as the stress conditions [61], can also be used to test our ideas, if the overall effect on protein localization of these events is known.

In conclusion, we have used a physical theory to estimate the order of magnitude of the inward and outward pressures originating in various mechanisms. In contrast to the widespread notion that the dominant contribution to the nuclear pressure results from the chromatin (which is widely believed to be the primary determinant of nuclear volume), our analysis suggests that localization of proteins and RNA molecules in the cytoplasm and nucleoplasm by active, nucleocytoplasmic transport is responsible for the dominant contributions to the inward and outward pressures, and hence determines nuclear volume. Motivated by this, we formulated a predictive model for the volume of the nucleus that is based on the osmotic pressure of the preferentially-localized, soluble molecules rather than the mechanical properties of large complexes such as the chromatin and cytoskeleton. While these structures may contribute important shear forces that modulate nuclear shape, they do not determine its volume. Our minimal model predicts that the ratio of the volumes of the nucleoplasm and cytoplasm is robust to a wide variety of biophysical and biochemical perturbations, an unexplained observation that is more than a century old. Beyond the prediction of the constant NC ratio, our work may impact the field of nuclear mechanobiology, since it highlights the potential role of osmotic pressures of the localized, soluble molecules and delineates the distinct response of the nucleus to shear forces and pressures. This also suggests that the mechanical response of the nucleus, which has so far been attributed to either the lamina or chromatin [49, 62], should be analyzed more carefully to acknowledge the contribution of osmotic pressures to nuclear mechanics.

## 5 Acknowledgements

We are grateful to Omar Adame Arana, Ram Adar, Gaurav Bajpai, Dennis Discher, Ming Guo, Hagen Hofmann, Jerome Irianto, Paul Janmey, Amit Kumar, Emmanuel Levy, Dana Lorber, Matthieu Piel, and Talila Volk for valuable discussions. We are especially grateful to Michael Elbaum for the critical reading of the manuscript and helpful comments. The research was supported by the Volkswagen foundation Grant, the Weizmann-Curie Grant, the Fern and Manfred Steinfeld Professorial Chair, Benoziyo Endowment Fund for Advancement of Science, Henry Krenter Institute for Biomedical Imaging and Genomics, Harold Perlman Family, and the Pearlman Grant.

