## Supplementary Information for "Balance of osmotic pressures determines the volume of the cell nucleus"

#### 1 Gibbs-Donnan calculation for the osmotic pressure of chromatin counterions

The limit we consider leads to an upper bound for the osmotic pressure of the chromatin counterions. In particular, we assume that the counterions are not localized near the chromatin fiber and their concentration is uniform within the nucleoplasm. Localization and inhomogeneity of the ion concentration lowers their contribution to the osmotic pressure. The concentrations of the different ions are then determined by two conditions: (1) Equality of the electrochemical potentials of the ions in the cytoplasm and nucleus. This potential, in the limit of a dilute solution, is the sum of entropic contribution equal to the log of the concentration of that ion species and a constant electrostatic term whose contribution is of opposite sign for positive and negative charges, $k_B T \log\left(c_i^{\pm} v_0\right) \pm e\nu\Phi_i$; where $k_B T$ is the thermal energy, $c_i^{\pm}$ is the concentration of positive/negative monovalent ions in compartment $i = c, n$ (cytoplasm or nucleoplasm), $v_0$ is the ion volume, taken to be the same for cations and anions, $e$ is the electron charge, $\nu$ is the valence of the ion, and $\Phi_i$ is the electrostatic potential in compartment $i$. Equality of the electrochemical potential is an equilibrium condition, which is appropriate for ions that can freely diffuse through the NE which has only passive channels connecting the nucleoplasm and cytoplasm and no ion pumps. (2) The nucleoplasm and the cytoplasm are both electro-neutral. The electrostatic energy cost of mesoscopically charged cytoplasm or nucleus is very large and thus suppresses such distributions [1].

In our minimal model for osmotic pressure of the counterions, the nucleoplasm comprises three types of solutes: (1) positive monovalent ions, (2) negative monovalent ions, and (3) chromatin; the cytoplasm, which lacks chromatin, contains only the ions. We denote the respective concentrations of positive and negative monovalent ions in the cytoplasm by $c_c^+$ and $c_c^-$ and their respective concentrations in the nucleoplasm by $c_n^+$ and $c_n^-$. The net, negative charge concentration of the chromatin (in units of electron charges) in the nucleoplasm is written as $c_{ch}^-$. From condition (1) of equality of electrochemical potentials for the two types of ions, the concentrations of the monovalent ions are related by $k_B T \log\left(c_c^+/c_n^+\right) = -k_B T \log\left(c_c^-/c_n^-\right)$. This relation is equivalent to the following, simpler relation, $c_c^+ c_c^- = c_n^+ c_n^-$. We note that because the chromatin cannot cross the NE, it does not reach electrochemical equilibrium and it does not satisfy these equations.

Condition (2) for the overall electro-neutrality of the cytoplasm and nucleoplasm, leads to the following two equations, $c_c^+ = c_c^-$ and $c_n^+ = c_n^- + c_{ch}^-$, for each compartment respectively. For simplicity, we work in the limit where the volume of the cytoplasm is much larger than that of the nucleus, so that ion exchange with the nucleus does not significantly change the ion concentration in the cytoplasm. This simplification allows us to approximate the cytoplasmic concentrations of the positive and negative ions as the intracellular salt concentration $c_c^+ = c_c^- \approx c_{salt}$; corrections to this approximation are not expected to change the order of magnitude of our pressure estimates. The equations $c_c^+ = c_c^- \approx c_{salt}$, $c_n^+ = c_n^- + c_{ch}^-$, and $c_c^+ c_c^- = c_n^+ c_n^-$ relate the concentrations of the monovalent ions in the nucleoplasm to the salt concentration and the concentration of chromatin counterions:

$$c_n^+ = \frac{\sqrt{\left(c_{ch}^-\right)^2 + 4c_{salt}^2} + c_{ch}^-}{2} \tag{1}$$

$$c_n^- = \frac{\sqrt{\left(c_{ch}^-\right)^2 + 4c_{salt}^2} - c_{ch}^-}{2} \tag{2}$$

The net contribution of the monovalent ions to the osmotic pressure is simply the difference between the total concentrations of all the ions in the nucleoplasm and all the ions in the cytoplasm, multiplied by $k_BT$:

$$p_c = k_BT\left(c_n^+ + c_n^- - 2c_{salt}\right) \approx k_BT\frac{\left(c_{ch}^-\right)^2}{4c_{salt}} \tag{3}$$

where the rightmost expression results from expansion of $\sqrt{\left(c_{ch}^-\right)^2 + 4c_{salt}^2}$ for the biological case that $c_{salt} \gg c_{ch}^-$ (see main text). Substituting the relation between the net chromatin charge concentration, the net chromatin charge, and the nuclear volume, $c_{ch}^- = N_{ch}/V_n$, into Eq. 3 results in Eq. 3 in the main text.

We note that our simple model of the nuclear electrostatics that considers only one type of positive, monovalent ions, one type of negative, monovalent ions, and chromatin, does not capture the true and varied composition of the cellular cytoplasm and nucleoplasm. The cytoplasm and nucleoplasm contain a multitude of ion species with various valencies, as well as charged proteins. However, the majority of the positive and negative ions in the cells are potassium ions and phosphate ions [2], respectively. This justifies our simplification that the positive and negative ions are all of the same type. Furthermore, while our choice of approximating all the ions as monovalent may affect the detailed, numerical value of the estimate, it is not expected to change its order of magnitude. In contrast, the cytoskeleton which is also negatively charged is not included in our estimate either. Its effect on the osmotic pressure can only lower our upper bound estimate. This is because the charge of the cytoskeleton is negative and confined to the cytoplasm, so that its contribution to the osmotic pressure difference between the cytoplasm and nucleoplasm only reduces our estimated contribution of the negatively charged chromatin that is confined to the nucleus. If the concentration of the cytoskeletal charge is comparable to that of the chromatin, it may significantly reduce the osmotic pressure difference. Therefore, the pressure predicted by 3 is an upper bound to the pressure contributed by the chromatin counterions.

### 2 Localized protein pressure

Study shows that in X. laevis oocytes, most of the cellular proteins are preferentially localized either to the cytoplasm or to the nucleoplasm [3]. Each of those proteins contributes to the net osmotic pressure of the compartment to which it is preferentially localized, a contribution proportional to its concentration in the compartment. This concentration is difficult to measure, since fractionation protocols of nucleoplasm and cytoplasm many times use detergents which perforate the NE, mixing the proteins of the nucleoplasm and cytoplasm [3]. For that reason, concentrations of cellular proteins are commonly measured and calculated in a cell-averaged manner, using total amount of proteins in the entire cell and the volume of the entire cell. The cell-averaged concentration of a localized protein is lower than its concentration in the compartment to which it is preferentially-localized and larger than its concentration in the other compartment. Therefore, the use of cell-averaged protein concentrations may potentially lead to underestimation of the net contribution of preferentially-localized proteins to the osmotic pressure. In this Appendix, we relate the compartment-specific osmotic pressures due to preferentially-localized proteins to the cell-averaged concentrations of those proteins for the important case discussed in the main text: The NE is relaxed and the osmotic pressure due to the localized proteins is the dominant contribution to the osmotic pressures of the two compartments.

In the cell, nucleocytoplasmic transport of proteins (and RNA molecules) through the nuclear pore complexes (NPCs) occurs via a shuttling mechanism that is coupled to existing concentration gradients of GDP-Ran and GTP-Ran across the NE [4, 5, 6]. These gradients are maintained by active enzymatic activity that differs in the cytoplasm and nucleoplasm. Due to the active nature of these enzymatic activities, the steady-state of the shuttling mechanism may be out-of-equilibrium, thus violating detailed balance in which the import and export rates are equal (e.g., due to diffusion that would occur for a passive channel). Thus, non-equilibrium activity implies that the import rate, $k_i$, is different than the export rate, $k_e$. In steady-state, in any time interval, the amount of any specific protein species that is imported must be equal to its amount that is exported. In the framework of first order kinetics, this condition relates the differing import and export rates to the cytoplasmic, $c_c$, and nucleoplasmic, $c_n$, protein concentrations: $k_i c_c = k_e c_n$. This implies that the ratio of the nucleoplasmic and cytoplasmic concentrations of the protein is equal to the ratio of the import and export rates, $c_n/c_c = k_i/k_e$. Since enzymatic transport activity can result in unequal import and export rates, the protein will be enriched in one compartment relative to the other.

Different protein species have different affinities to the molecular components involved in the nucleoplasmic shuttling mechanism, which suggests that each species that is actively transported may have different import and export rates and thus a different relative enrichment in one of the two compartments [5, 6]. We denote the relative enrichment of protein $i$ by $\kappa_i$, defined as the ratio of the concentrations of the protein in the nucleoplasm and in the cytoplasm, $\kappa_i \equiv c_{i,n}/c_{i,c}$; if $\kappa_i > 1$ then protein $i$ is actively transported and preferentially-localized to the nucleus, if $\kappa_i < 1$, then protein $i$ is actively transported and preferentially-localized to the cytoplasm. For simplicity, we consider here only proteins that are actively transported to one of the compartments so that $\kappa_i, \kappa_j \neq 1$ for any protein in the summations below.

Using the definition of the relative enrichment, we write the net contribution of the preferentially-localized proteins to the osmotic pressures of the cytoplasm or the nucleoplasm as $p_c$ and $p_n$, respectively; these are related to the concentrations of the proteins and their relative enrichment by:

$$p_c = k_B T \sum_i (c_{i,c} - c_{i,n}) = k_B T \sum_i (1 - \kappa_i)\, c_{i,c} \tag{4}$$

$$p_n = k_B T \sum_j (c_{j,n} - c_{j,c}) = k_B T \sum_j (\kappa_j - 1)\, c_{j,c} \tag{5}$$

where the subscript $i$ denotes summation over different protein species preferentially-localized to the cytoplasm and the subscript $j$ denotes summation over different protein species preferentially-localized to the nucleoplasm. We denote the total cell-averaged concentrations of all the proteins that are preferentially-localized to the cytoplasm and all the proteins that are preferentially-localized to the nucleoplasm as $\bar{c}_c$ and $\bar{c}_n$, respectively. These total, cell-averaged concentrations are expressed using their compartment-specific concentrations and the free volumes of the compartments as follows:

$$\bar{c}_c = \frac{\sum_i \left( (V_c - V_{c,m})\, c_{i,c} + (V_n - V_{n,m})\, c_{i,n} \right)}{(V_c - V_{c,m}) + (V_n - V_{n,m})} = \frac{\sum_i (\alpha c_{i,c} + c_{i,n})}{\alpha + 1} = \frac{\sum_i (\alpha + \kappa_i)\, c_{i,c}}{\alpha + 1} \tag{6}$$

$$\bar{c}_n = \frac{\sum_j \left( (V_c - V_{c,m})\, c_{j,c} + (V_n - V_{n,m})\, c_{j,n} \right)}{(V_c - V_{c,m}) + (V_n - V_{n,m})} = \frac{\sum_j (\alpha c_{j,c} + c_{j,n})}{\alpha + 1} = \frac{\sum_j (\alpha + \kappa_j)\, c_{j,c}}{\alpha + 1} \tag{7}$$

where as before, subscripts $i$ and $j$ respectively denote summation over protein species that are preferentially-localized to the cytoplasm and nucleoplasm. The free volumes of the cytoplasm and nucleoplasm are respectively denoted as $V_c - V_{c,min}$ and $V_n - V_{n,min}$, and $\alpha$ denotes their ratio, $\alpha \equiv (V_c - V_{c,m}) / (V_n - V_{n,m})$. Using Eqs. 6 and 7, we write the total, cell-averaged concentration of all localized proteins (cytoplasm and nucleoplasm) as

$$\bar{c}_c + \bar{c}_n = \frac{\sum_i (\kappa_i - 1)\, c_{i,c} + \sum_j (\kappa_j - 1)\, c_{j,c}}{\alpha + 1} + \sum_i c_{i,c} + \sum_j c_{j,c} \tag{8}$$

For the case we consider in this Appendix, that the NE is relaxed and the dominant contributions to the inward and outward pressures originate from localization of proteins, the osmotic pressures of the preferentially-localized proteins in the two compartments, $p_c$ and $p_n$, are equal due to mechanical equilibrium (see main text). Using Eqs. 4 and 5, and the definition of the relative enrichments, the equality of the two pressures results in the following two relations:

$$\sum_i (\kappa_i - 1) c_{i,c} = -\sum_j (\kappa_j - 1) c_{j,c} \quad (9)$$

$$\sum_i c_{i,n} + \sum_j c_{j,n} = \sum_i c_{i,c} + \sum_j c_{j,c} \quad (10)$$

When Eq. 9 is substituted into Eq. 8 for the total average, cellular concentrations of all localized proteins (cytoplasm and nucleoplasm), we find that $\bar{c}_c + \bar{c}_n = \sum_j c_{j,c} + \sum_i c_{i,c}$. This indicates that the total cell-averaged concentrations of all proteins that are actively transported, is equal to the sum of the cytoplasmic concentrations of the proteins that are preferentially-localized to the cytoplasm, and the cytoplasmic concentrations of the proteins that are preferentially-localized to the nucleoplasm. In turn, Eq. 10 indicates that this is also equal to the sum of the nucleoplasmic concentrations of the proteins that are preferentially-localized to the nucleoplasm and the nucleoplasmic concentrations of the proteins that are preferentially-localized to the cytoplasm. This conclusion is also true for the simple case considered in the main text, where there are $N_c^l$ proteins that are completely (rather than preferentially) localized to the cytoplasm and $N_n^l$ that are completely localized to the nucleoplasm. In this case our conclusion indicates that from the equality of osmotic pressures of these proteins (for the case that the NE is relaxed), $N_c^l / (V_c - V_{c,m}) = N_n^l / (V_n - V_{n,m})$, the following two identities follow, $\left(N_c^l + N_n^l\right) / (V_c - V_{c,m} + V_n - V_{n,m}) = N_n^l / (V_n - V_{n,m})$ and $\left(N_c^l + N_n^l\right) / (V_c - V_{c,m} + V_n - V_{n,m}) = N_c^l / (V_c - V_{c,m})$. These identities are proven by the following simple algebraic steps:

$$\begin{aligned}
\frac{N_c^l}{(V_c - V_{c,m})} &= \frac{N_n^l}{(V_n - V_{n,m})} \Rightarrow \\
\frac{N_c^l}{N_n^l} &= \frac{(V_c - V_{c,m})}{(V_n - V_{n,m})} \Rightarrow \\
\frac{N_c^l + N_n^l}{N_n^l} &= \frac{(V_c - V_{c,m}) + (V_n - V_{n,m})}{(V_n - V_{n,m})} \Rightarrow \\
\frac{N_c^l + N_n^l}{(V_c - V_{c,m}) + (V_n - V_{n,m})} &= \frac{N_n^l}{(V_n - V_{n,m})} = \frac{N_c^l}{(V_c - V_{c,m})}
\end{aligned}$$

This calculation indicates that if about 80% of the proteins are preferentially-localized to either compartment [3], and the total osmotic pressure of all proteins is about 10 kPa [7], then the osmotic pressures in the cytoplasm and nucleoplasm due to localization of proteins is of the order of 8 kPa. As explained in detail in the main text, this 8 kPa estimate is an upper bound for the contribution of the localized proteins to the inward and outward pressures.

### 3 Relation to nucleocytoplasmic transport

In this appendix, we generalize the values used in the main text, $N_c^l$ and $N_n^l$, which represent the number of solutes completely localized to the cytoplasm and nucleoplasm, respectively. The main text focuses on the simple case that there is only one species of macromolecular solute that is found only in the cytoplasm and another species of solute that is found only in the nucleoplasm. Here, we generalize these values to the biologically realistic scenario where there are multiple macromolecular solutes (proteins and RNA molecules) which are preferentially-localized rather than completely-localized. Namely, they are found in both the cytoplasm and nucleoplasm, although in different concentrations, which are determined by the characteristics of the active nucleocytoplasmic transport mechanisms. As explained in the section "Localized protein pressure" above, the nucleoplasmic and cytoplasmic concentrations of each protein $i$ that is actively transported are different. This is quantified by the relative enrichment $\kappa_i$, defined as the ratio between the cytoplasmic and nucleoplasmic concentrations of the protein $\kappa_i \equiv c_{i,n}/c_{i,c}$; if $\kappa_i > 1$ then solute $i$ is actively transported to and is preferentially localized to the nucleus, if $\kappa_i < 1$, then solute $i$ is actively transported to and is preferentially localized to the cytoplasm. Here, we consider only solutes that are actively transported to one of the compartments so that $k_i \neq 1$ for any solute $i$ in the summations below.

In the case that the NE is relaxed, mechanical balance is characterized by Eq. 1 in the main text with $\sigma = 0$, which indicates that the osmotic pressure of all the proteins localized on each of the two sides of the NE is equal. Since the small solutes that are not transported have the same concentration in the nucleoplasm and cytoplasm, the contribution of each small solute species to the net osmotic pressures of either compartment is zero. The osmotic pressure balance of the remaining large proteins that are actively transported is written in the approximation of an ideal solution as:

$$k_B T \sum_i c_{i,n} = k_B T \sum_i c_{i,c} \tag{11}$$

where the summation is over all of the proteins that are actively transported so that their concentrations in the cytoplasm and nucleoplasm are unequal.

In addition to Eq. 11, the number of molecules of each protein species is conserved, so that $c_{i,c}$ and $c_{i,n}$ satisfy the following relation

$$N_i = (V_c - V_{c,m})\, c_{i,c} + (V_n - V_{n,m})\, c_{i,n} \tag{12}$$

where $N_i$ is the total, cellular copy number of protein $i$, and $V_c - V_{c,m}$ and $V_n - V_{n,m}$ are the respective free volumes of the cytoplasm and nucleoplasm (see main text). We note that over long enough timescales, proteins are produced and degraded in vivo, thus our use of the conservation of proteins is only appropriate when the experimental timescales are much shorter than the protein production and degradation timescales (which are of the order of 10 hours [2, 8]).

Eq. 12, together with the definition of the relative enrichment, $\kappa_i \equiv c_{i,n}/c_{i,c}$, can be used to express the concentrations of the proteins in the cytoplasm and nucleoplasm using the free volumes of the two compartments and the total, cellular copy numbers of each species of protein:

$$c_{i,c} = \frac{N_i}{(V_c - V_{c,m}) + \kappa_i (V_n - V_{n,m})} \tag{13}$$

$$c_{i,n} = \frac{\kappa_i N_i}{(V_c - V_{c,m}) + \kappa_i (V_n - V_{n,m})} \tag{14}$$

Substituting Eqs. 13 and 14 into the condition for the mechanical equilibrium of the relaxed NE, Eq. 11 and multiplying by $V_n - V_{n,m}$, results in an implicit equation for the ratio of the free volumes $(V_c - V_{c,m}) / (V_n - V_{n,m})$

$$\sum_i \frac{\kappa_i N_i}{\frac{(V_c - V_{c,m})}{(V_n - V_{n,m})} + \kappa_i} = \sum_i \frac{N_i}{\frac{(V_c - V_{c,m})}{(V_n - V_{n,m})} + \kappa_i} \tag{15}$$

Solving Eq. 15 to find the ratio between the free volumes $(V_c - V_{c,m}) / (V_n - V_{n,m})$ as an explicit function of the different copy numbers, $N_i$ and the localization ratios, $\kappa_i$, is not possible. However, this equation implies that this ratio is a constant that is related to the $N_i$-s and the $\kappa_i$-s. Furthermore, Eq. 15 allows us to derive an important identity that relates the ratio of the free volumes to the total number of proteins in each compartment (those that are preferentially localized to the compartment and other species in that compartment that are preferentially localized to the other compartment), $N_c^l$ and $N_n^l$; notably, those correspond to the quantities used in the minimal model presented in the main text. To do this, we first express $N_c^l$ and $N_n^l$ as the following functions of the free volumes, protein copy numbers $N_i$, and relative enrichments $\kappa_i$:

$$N_c^l = \sum_i (V_c - V_{c,m})\, c_{i,c} = \sum_i \frac{(V_c - V_{c,m})\, N_i}{(V_c - V_{c,m}) + \kappa_i\,(V_n - V_{n,m})} \tag{16}$$

$$N_n^l = \sum_i (V_n - V_{n,m})\, c_{i,n} = \sum_i \frac{\kappa_i\,(V_n - V_{n,m})\, N_i}{(V_c - V_{c,m}) + \kappa_i\,(V_n - V_{n,m})} \tag{17}$$

Multiplication of Eq. 15 by $(V_c - V_{c,m}) \,/\, (V_n - V_{n,m})$, followed by substitution of Eqs. 16 and 17 into the resulting product, results in the relation

$$\frac{V_c - V_{c,m}}{V_n - V_{n,m}} = \frac{N_c^l}{N_n^l} \tag{18}$$

This is the same conclusion that we arrived at in the main text, where we used $N_c^l$ and $N_n^l$ to denote a simple case where only one species is completely localized in each compartment. Therefore, the detailed calculations presented in this appendix justify the use of $N_c^l$ and $N_n^l$ in the minimal model presented in the main text. The rest of the results presented in the main text all rely on Eq. 18 which we derived here rigorously, and are thus equally justified.

### 4 Cytoplasmic and nuclear volumes in the non-ideal limit

In the main text, we solved Eqs. 4 and 7 that represent mechanical equilibrium across the NE and the plasma membrane, for the ideal solution limit where the minimal volumes of the cytoplasm and nucleoplasm are negligible compared to their respective volumes. This yielded expressions 9 and 10 in the main text that relate the volumes of the cytoplasm and nucleoplasm. These equations indicated that the ratio between the volume of the nucleoplasm and the cytoplasm (NC ratio) depends only on the ratio of the numbers of completely-localized proteins in the nucleoplasm and in the cytoplasm. However, the corrections to the ideal limit which we used to determine the volumes of the cytoplasm and nucleoplasm may be significant, since the minimal volume of the cell, and presumably the minimal volumes of the separate compartments, may reach tens of percent of their total volumes [9]. In this Appendix, we now include the effects of the minimal volumes (steric, excluded volume of the various solutes). We show that the prediction regarding the NC ratio for $\sigma = 0$ is also valid in the non-ideal limit, as long as the volumes of the non-diffusive complexes in the compartments are small compared with the total volumes of the compartments.

In the non-ideal limit, the free volumes of the two compartments, $V_c - V_{c,m}$ and $V_n - V_{n,m}$, replace the total volumes of the compartments in Eqs. 9 and 10 in the main text, which results in the following equations

$$V_c - V_{c,m} = \frac{N_c^l}{C}\left(1 + \frac{N}{N^l}\right) \tag{19}$$

$$V_n - V_{n,m} = \frac{N_n^l}{C}\left(1 + \frac{N}{N^l}\right) \tag{20}$$

where $N$ is the total number of non-localized, small solutes in the cell, $N_c^l$ and $N_n^l$ are the number of solutes that are completely-localized to the cytoplasm and nucleoplasm, respectively, $C$ is the concentration of solutes in the extra-cellular environment, and $N^l = N_c^l + N_n^l$ is the total number of localized solutes in the entire cell.

In experiments, the total volumes of the two compartments, $V_n$ and $V_c$, are measured rather than their free volumes, $V_n - V_{n,m}$ and $V_c - V_{c,m}$. This motivates us to express the minimal volumes of the two compartments using the numbers of different solutes defined above, in order to rewrite Eqs. 19 and 20 as explicit functions of the observables, $V_c$ and $V_n$. The minimal volumes of the two compartments are the total volume of all their non-solvent components, including small, non-localized solutes, large, localized solutes, and non-diffusive structures within each the two compartments (e.g. cytoskeleton and chromatin, respectively). To account in an algebraically simple manner for the different sizes of localized vs. non-localized solutes [10] in our minimal model, we consider different average volumes for these two types of molecules: The volumes of the various small, non-localized solutes (including ions and small molecules) are taken to be equal, $v_s$, while the volumes of the various, large, localized solutes are also taken to be equal, $v_\ell$, where $v_\ell > v_s$. The average volumes of the different solutes allows us to write the minimal volumes as $V_{c,m} \approx v_s N_c + v_\ell N_c^l + V_{c,s}$ and $V_{n,m} \approx v_s N_n + v_\ell N_n^l + V_{n,s}$, where respectively: $N_c$ and $N_n$ are the numbers of non-localized solutes in the cytoplasm and nucleoplasm, $N_c^l$ and $N_n^l$ are the total numbers of localized solutes in the cytoplasm and nucleoplasm, and $V_{c,s}$ and $V_{n,s}$ are the volumes of the non-diffusive structures in the cytoplasm and nucleoplasm. Substituting the expressions for the minimal volumes and the ratio of the free volumes (Eq. 8 in the main text) into main-text Eqs. 5 and 6 for $N_c$ and $N_n$, and the resulting expressions into Eqs. 19 and 20, we derive the following expressions for the cytoplasmic and nuclear volumes, $V_c$ and $V_n$

$$V_c = N_c^l \left(1 + \frac{N}{N^l}\right)\left(\frac{1}{C} + \bar{v}\right) + V_{c,s} \tag{21}$$

$$V_n = N_n^l \left(1 + \frac{N}{N^l}\right)\left(\frac{1}{C} + \bar{v}\right) + V_{n,s} \tag{22}$$

where $N$ is the total number of non-localized solutes in the cell, $N^l = N_c^l + N_n^l$ is the total number of localized solutes in the cell, and $\bar{v} = \left(v_s N + v_\ell N^l\right) / \left(N + N^l\right)$ is the average volume of a solute in the cell. We observe that if $V_{c,s} \ll V_c$ and $V_{n,s} \ll V_n$, the ratio of the nuclear and cellular volumes is well-approximated by the ideal solution limit (see main text) and is again equal to the ratio, $N_n^l/N_c^l$, of the numbers of their respectively localized proteins. This is the typical biological situation in which the total volumes of the non-diffusing structures in the cytoplasm and nucleoplasm, $V_{c,s}$ and $V_{n,s}$, are much smaller than the total volumes of their respective compartments. The volume of the cytoskeleton, which is presumably the largest non-diffusing structure in the cytoplasm, is only few percent of the volume of the cytoplasm [11]. Similarly, the calculations below of the total volume fraction of (bare) chromatin in human and S. pombe nuclei, based on the structural properties of DNA and histone proteins, show that it is of the order of one percent of the volume of the nucleoplasm.

The volume of chromatin in the nucleus is the sum of the total volumes of the DNA and its associated proteins. Since the most abundant and largest DNA-binding, protein complexes are the histones, the total volume of the chromatin is well-approximated by the sum of the DNA and histone volumes. From crystallographic structural measurements of DNA and histone octamers, the physical dimensions of the two biomolecules are known: DNA occupies a volume approximated by a cylinder of radius of 1 nanometer whose contour length is 0.34 nm multiplied by the number of base pairs [12]. This results in a DNA volume which is about 1 nm$^3$ per base pair. A histone octamer occupies a volume approximated by a cylinder whose diameter is 6.5 nm and height 6 nm [12], so that the volume of each histone octamer is $\sim$200 nm$^3$. The average density of histones in a genome is typically one histone octamer per $\sim$200 base pairs [12], which indicates that the presence of histones contributes about 1 nm$^3$ per base pair of DNA, similar to the contribution of the DNA itself. Therefore, the total volume of the chromatin (in nm$^3$) of a eukaryotic organism is about twice the size of its genome in base pairs.

In human cells, the diploid genome is $6.4 \cdot 10^9$ base pairs long [2], which implies that the total volume of the chromatin is $\sim 12.8 \cdot 10^9$ nm$^3$. In S. pombe cells, the diploid genome is $27.6 \cdot 10^6$ base pairs long [13], which similarly implies that the total volume of the chromatin is $\sim 55.2 \cdot 10^6$ nm$^3$. In order to calculate the volume fraction of chromatin in the nuclei of the two organisms, we divide the chromatin volumes of human and S. pombe cells by the respective volumes of their typical nuclei, $\sim$ 1000µm$^3$ [9] and $\sim$ 17µm$^3$ [14]. We find that the volume fractions of chromatin in the two species is $\sim$ 1.3% for humans and $\sim$ 0.32% for S. pombe, both are of the order of one percent or less. Of course this does not include the water of hydration, screening cloud of counterions or other proteins that might bind to chromatin.

In conclusion, the low volume fractions of the cytoskeleton and chromatin, presumably the largest non-diffusive structures in the cell, allows us to approximate the NC ratio calculated from Eqs. 21 and 22 in the non-ideal solution limit, in a manner similar to the one calculated in the ideal limit that is presented in the main text.
